# Sex-specific perching: Monitoring of artificial plants reveals dynamic female-biased perching behavior in the black soldier fly, *Hermetia illucens* (Diptera: Stratiomyidae)

**DOI:** 10.1101/2024.07.23.604854

**Authors:** Noah B Lemke, Lisa N Rollinson, Jeffery K Tomberlin

## Abstract

Artificial perches are implemented by many companies that mass-rear the black soldier fly, *Hermetia illucens,* to create a more natural breeding environment or provide additional surface area for flies to rest. However, basic information related to perching behavior is lacking. This experiment tested the effect of adding perches to breeding cages on fitness and behavior. Perches were constructed from artificial leaves affixed to wooden dowels inserted into foam blocks, placed in the center of cage floors. The four treatment-levels had an added surface area of 0.00-, 0.04-, 0.26-, and 0.34-m^2^. Each 0.93-m^3^ cage was supplied with 90 male and 90 female flies, and female thoraxes marked with acrylic paint. Beyond the tested range, a linear model suggests that 1.00-m^2^ additional surface area can accommodate a 1.46-fold increase in flies without negative fitness impacts. Time-series analysis revealed; (a) females utilized perches 1.42-times more often than males across two trials; (b) especially in the morning where the difference could be as high as 2.56-times more than males; (c) this decreased to 0.20-1.57 times more females than males by 1600 h, and (d) this cyclical pattern repeated each day throughout the week with decreasing female-bias, starting from 2.41-times more females on Day 1 to 0.88-1.98-times more females than males on Day 6. These dynamics are likely due to the presence of male flies engaging in aerial contests near lamps providing light needed for mating, especially during early hours and early adulthood, aligning with prior knowledge of black soldier fly mating behavior.

**LAY SUMMARY:** Female black soldier flies perch on artificial plants much more often than males, especially early in the day and early in their reproductive lives, times when males are competing with one another in aerial swarms.

## Introduction

The study of habitat structural complexity has been a longstanding and pervasive theme in ecology (Loke and Chisholm 2022), dating back to the 1950’s and before with Huffaker’s famous manipulation experiment in which he alternated the arrangement of an orange and rubber ball ‘universe’, affecting the population dynamics of herbivorous and predatory mites (Acaridae) (Huffaker 1958). For decades, researchers have endeavored to characterize both biotic and biotic factors that impact animal behavior (Hastings et al. 2007).

For neotropical (Kaya and Gaugler 1993) detritovores like the black soldier fly, *Hermetia illucens* (L. 1758) (Diptera: Stratiomyidae) (Lemke et al. 2023), Jenny’s State Equation (Jenny 1941), can be used as a formalized way to link spatially-explicit factors in a single conceptual framework. The model posits that temperature (Chia et al. 2018) and humidity (Holmes et al. 2012; Bekker et al. 2021) increase the rate of nutrient cycling and soil formation, which is also dependent on several factors such as the nutritional quality of the substrate (Barragan-Fonseca et al. 2019; Barragan-Fonseca et al. 2021) as well as larval competition (Jones and Tomberlin 2019).

The arrangement of objects in space is also linked *de facto* to selection and resource partitioning (Bell et al. 1991). For this reason, it is likely that spatial complexity affects the efficiency of larvae as they process waste, such as through substrate depth (Brits 2017), particle size (Brits 2017), and heterogeneity of non-digestible objects (Shishkov et al. 2019; Liu et al. 2022). Although spatial factors such as these are understood to affect the growth and development of black soldier fly larvae, the effect of habitat conformation on reproductive behavior is less appreciated, even though it is known that the alteration of habitat (via domestication) can directly impact adult behavior and ultimately fitness (Price 1984).

Altering habitat structure (whether through natural or artificial processes) can impact species behavior across multiple scales. Knowledge of this bottom-up process has been used in various applied contexts ranging from the design of artificial reefs (Bohnsack 1991) and urban greenspaces (Golstein-Golding 1991) to nature preserves (Usher 1991). As a specific example, in the mass-reared *Ceratitits capitata* (Weidemann, 1824) (Diptera: Tephritidae), modifying enclosures with slats (Liedo et al. 2007) was shown to increase the number of lekking sites (Bradbury et al. 1986; Alcock 1987; Höglund and Alatalo 1995; Choe and Crespi 1997), which enhanced reproductive performance by reducing male-aggression (Briceño et al. 1999).

Yet, for the similarly lekking black soldier fly (Tomberlin and Sheppard 2001; Birrell 2018), little is known about how adults explicitly interact with the structure of their habitat in any context. Besides a singular report (Tomberlin and Sheppard 2001), in which the authors describe males (with 91.9% (n=109) male-to-female sex ratio) perching on Kudzu *Pueraria montana* (Loer) (Merr.) (Tomberlin and Sheppard 2001) nothing is known about adult behavior in the wild.

Colony production methods for the black soldier fly initially describe providing artificial plants (50-cm globes, with leaves that resemble ivy measuring 3.8-7.6 cm) to flies that were misted twice-daily with an automatic water sprayer (Sheppard et al. 2002). Because of this, or simply out of intuition, many industrial producers supply their cages with artificial plants (Isa and Hasan 2021), apparently to provide resting area (Isa and Hasan 2021), or increase the absolute surface area available relative to each fly (Meneguz et al. 2023). However, the direct effects on fecundity or fitness have not been quantified, nor any on behavior (i.e., perching) or insect welfare (Boppré and Vane-Wright 2019; Barrett et al. 2022) (e.g., stress-reduction (Cattaneo et al. 2024)). If the inclusion of perches has no effect on fitness, this potentially represents an unneeded cost to cages used in a mass-rearing facility.

Perching behavior in insects is theorized to aid with mate-locating (Scott 1974; Alcock 1984), thermoregulation (Young 1984), and the reduction of intra-specific competition (Noriega and Vulinec 2021) by allowing insects to vertically stratify. With regards to mate-locating behaviors, insects often perch as they wait for conspecifics to approach (Tomberlin and Sheppard 2001), meaning that theoretically the inclusion of perches could facilitate reproduction by promoting natural behaviors.

Using the McCoy and Bell’s conceptual model (1991), habitat structural complexity was defined as the arrangement of objects in the environment, that possess a functional interaction and the organism being studied, at a suitable scale, but which are not explicitly necessary for survival, or incompatible with life. Critically, McCoy and Bell’s definition (1991) removes from consideration the oviposition substrate, and possibly the cage itself, since the enclosure might inhibit processes that would happen if flies were uncaged. This perspective hence renders artificial plants as the objects of focus. Such a functional interaction could be that plants or similar structures are utilized by males to first establish lekking sites (Scott 1974; Alcock 1984), where females will visit to mate before returning to the oviposition site (Höglund and Alatalo 1995; Tomberlin and Sheppard 2001; Lemke et al. 2023).

We then ask whether an increasing gradient of habitat structural complexity likewise positively affects reproductive outcomes in black soldier flies, hypothesizing that there will be an indirect effect on fertile egg production, but this effect will diminish if complexity is too great with respect to scale. Furthermore, on the basis that males were previously observed to associate with plants (Tomberlin and Sheppard 2001), we expect that perching behavior will be primarily associated with males.

## Methods

We conducted all experimentation in the Forensic Laboratory for Investigative Entomological Sciences (FLIES) Facility (Texas A&M University, College Station, TX, USA). We conducted two trials. We modified methods from the following studies (Jones and Tomberlin 2021; Dickerson et al. 2024).

### Experimental Design

In keeping with prior studies (Jones and Tomberlin 2021; Laursen et al. 2022; Dickerson et al. 2024; Lemke 2024), we utilized twelve Insect-A-Hide Pop-Up Shelters (84 × 84 × 133 cm, L × W × H) (Lee Valley Tools Ltd., Ottawa, Ontario, Canada) as mating cages, and arranged them in a 2 × 6 array within room 9 of building 1043 of the F.L.I.E.S. Facility. We assigned treatments (CTRL, LOW, MED, HIGH) randomly, such that each treatment category had three replicates. To each cage, we fixed a 50-watt light-emitting diode (LED) (HK SPR AGTECH Trading LTD, Hong Kong, China) at a height of 101 cm and distance 1 cm from the cage door (Dickerson et al. 2024). During experimentation, we ran lights on a 12:12 L:D (6:00-18:00 h) to approximate both duration of natural light in a previous study (Lemke 2024) and in the tropics (personal observation) to which black soldier flies are native (Kaya et al. 2021). Because temperature and humidity were not controlled, we recorded these variables synchronously using a HOBO^®^ data logger MX1104 (Onset Computer Co., MA, USA).

### Artificial Plants

To construct artificial plants (See supplementary info for image), we used unfinished dowel rods (3.18mm × 15 cm: dia × L) (EBOOT, Guangdong, China) as stems and attached them to the underside of plastic artificial *Monstera deliciosa* Liebm. (Alismatales: Araceae) leaves (KUUQA, China) using strips of 18 mm wide masking tape (Scotch TM, St. Paul, MN, USA). The artificial leaves measured 12.2 cm (small) or 17.5 cm (medium) in length from base to tip and 11.0 cm and 17.1 cm from margin to margin, respectively. We elected not to use larger leaves from the same manufacturer because they were not rigid enough. We then affixed the dowel-rod stems (with leaves-attached) in a radial-fashion to phenolic floral foam blocks (3.81 × 7.62 cm, H x rad) (Decorat LTD, Mexico) with approximately even spacing in-between each stem. The respective number of leaves used to form LOW, MED, and HIGH treatment combinations are presented in Table 1.

**Table 1.** Treatment levels.

| **Table 1.** Treatment levels. |  |  |
| --- | --- | --- |
| **Level** | **Construction** | **Estimated Adaxial Leaf Area (m<sup>2</sup>)** |
| CTRL | Inverted pot only | 0.000000 |
| LOW | Pot + ½ small block + ½ medium block | 0.040061 |
| MED | Pot + 1 small block + 1 medium block | 0.260011 |
| HIGH | Pot + 2 small blocks + 2 medium blocks | 0.340133 |

We then stacked the floral foam ‘blocks’ (with leaves inserted by their stems) to form the respective treatment levels. Three small or three medium-sized leaves inserted into a floral foam block constituted a ‘half-block’, and six-leaves constituted a ‘full-block’. We offset the arrangement of blocks on the stack relative to one another to minimize the amount in which leaves overlapped spatially. In addition, we always placed blocks of small leaves above medium on the stack to mimic “new growth” of the fake plant. We secured the entire stack by plunging a singular dowel rod vertically through the core of the blocks. We placed the arrangements atop inverted nursery pots (15.25 cm D, 1.89 L) (Oubest, Fuzhou City, China), to mimic a potted plant. Controls only had inverted nursery pots.

### Leaf Area

We calculated an estimate for total added surface area, with respect to cage volume (0.93 m^3^) by importing photographs and reference scale from the manufacturer’s e-commerce website (Amazon.com) into ImageJ (Version 1.5h, BSD-2 Public License, NIH). Using the GUI-inbuilt measuring tool, we measured leaf lengths from the tip to the petiole along the midrib. We measured widths from the widest portions of the margins, perpendicular to the midrib. The top (adaxial) surface area we then approximated as the simple multiple of these two linear dimensions without subtracting any negative space for scalloped edges or lobed leaves. To net total added area, this dimension could be doubled; however, flies almost never utilized the bottom (abaxial) surface of the leaf to perch. Similarly, the Neotropical dung beetle *Canthon septemmaculatus* (Larr.) (Coleoptera: Scarabaeidae) only utilizes the top surface of leaves, albeit for thermoregulation (Young 1984).

### Rearing

We reared adult flies from 7-d-old larvae acquired from EVO Conversion Systems LLC (College Station, TX, USA) (Tomberlin et al. 2021) according to the methods of Lemke (2024, Chapter 3), with the following modifications:

(a) The first allotment of Gainesville diet (Hogsette 1992) (Producers Cooperative Association, Bryan, TX, USA) provided to the larvae weighed 5 kg, and the second weighed 3 kg, rather than an even 4 kg-4 kg split.
(b) During the grow-out periods, the walk-in incubator (although set to 26°C) experienced markedly different conditions (Trial 1: 23.22±2.00°C, 71.71± 9.78% RH; Trial 2: 25.67±1.52°C, 53.47±9.66% RH). This information we collected using a HOBO^®^ data logger MX1104 (Onset Computer Co., MA, USA). The second trial overlapped with an abnormal winter climactic event (Jan 12-16; Feb 10-11), during which sub-zero temperatures affected most of the continental United States.
(c) In addition, after sexing flies according to external genital anatomy (Munsch-Masset et al. 2023) and temporarily placing flies into BugDorm-1 holding cages (30 × 30 × 30 cm) (MegaView Science Co., Ltd, Taichung, Taiwan), the thoraxes of females only received a single dot of acrylic paint from a 3-mm tip Garde’n’ Craft Fine Point Marker (Uchida of America Corp, Torrance, CA, USA) (Jones and Tomberlin 2020; Muraro et al. 2024). Prior validation experiments show no negative effect against fecundity or fitness of painting females individually in this manner (Jones and Tomberlin 2020). Handling of flies prior to experimentation also has no reported detrimental effect on fitness (Tomberlin et al. 2009).

### Sequence of Experiment

On the evening prior to the start of each trial, we stocked each 0.93 m^3^ cage with 90 unpainted males and 90 painted females. At the top of the hour, from 0700-1800 h, we made behavioral observations (i.e., counts of mating events, count of ovipositing females in traps, counts of males and females occupying perches within each cage) visiting cages in a random order and alternating between two observers for 6-7-d. After this duration, the Pareto distribution for mating and oviposition events plateaued, which indicated that little additional behavioral information would be collected. This resulted in a minimum of 72 observations per trial that would then be averaged by day or by hour during statistical analyses. We misted each cage three times daily with 100 ml of reverse osmosis (RO) water (0700h, 1200 h, and 1800 h).

One the day following peak mating (i.e., highest total across all cages combined), which indicated the initiation of the post-mating interval, we added an attractant box and egg trap to each cage (Dickerson et al. 2024). This set-up consisted of plastic shoeboxes containing 1 kg of 70% moisture Gainesville diet inoculated with an aliquot of ∼1000 7-d-old larvae. We modified lids to allow volatiles to escape through a 12.7 (L) × 5 (W) cm rectangular slot, which was replaced with nylon screening. We constructed egg traps by taping together three strips of corrugated cardboard, measuring 10 (L) × 3.5 (W) × 1.25 (H) cm and placed them atop the lids. Previous work indicated that delaying substrate provisioning can have a positive effect on trapped eggs in conjunction with restricted cohort age (Lemke 2024).

We replaced cardboard egg traps daily at 12:00 h (Dickerson et al. 2024). We then harvested eggs following the methods of Lemke (2024). In addition to total weight of eggs per trap per cage per day, we also recorded the number of clutches in each trap to potentially better indicate the number of oviposition events that had occurred, as previous work also tended to show a lack of correlation between oviposition counts and egg weight (Lemke 2024). Any collected eggs we then placed in 30-ML solo cups and glass canning jars following the procedure from (Laursen et al. 2022; Dickerson et al. 2024) and incubated them within the same walk-in chamber (Trial 1: 27.19 ± 1.21 °C, 50.04 ± 7.33% RH; Trial 2: 25.59 ± 1.18 °C; 51.22 ± 5.57% RH). We monitored all collected eggs thereafter for five days (Lemke 2024) for hatch rate (Laursen et al. 2022; Dickerson et al. 2024).

### Non-Parametric Statistics

We performed all statistical analyses in RStudio (version 4.1.2) using tidyverse packages (Wickham et al. 2019). Visual inspection of data followed by Shapiro-Wilk test for Normality confirmed that the distribution of data (mating events, oviposition events, number of clutches, clutch weight, egg weight, and percent hatch) were non-normal (in all cases being highly skewed or zero-inflated). Therefore, we used nonparametric analyses (i.e., Kruskal Wallis *H*) to test whether treatment or increasing leaf area influenced fitness metrics.

### Perch Disparity Metric

Because painting females revealed obvious sex-specific behavioral patterns during experimentation, it was necessary to develop an index to describe perching patterns with respect to sex, which we call perch disparity.

Here defined, perch disparity is the simple difference of the count of females perching (per cage/per day) minus the count of males perching (per cage/per day); where values greater than 1 indicate a female bias, 0 indicates parity, and negative values indicate a male bias. Values for perch disparity were automatically calculated in Excel (version 2403, Build 17425.20176) by subtracting column totals from one another.

We cleaned, plotted, and fitted a curve to perching data using MS Excel’s built-in graphing tools. We did not transform data. We selected regressions based on the equation that maximized the R2-value while using the least number of terms. If this could not be achieved, then we chose the simplest expression. When calculating the area between two curves, we verified the difference of the two integrals using Wolfram Alpha (Wolfram Alpha LLC, Champaign, IL, USA).

### Time Series

To conduct a time series analysis, we imputed missing perch disparity data from unobserved time periods in MS Excel by calculating a 13 h moving average. We then superimposed the imputed values onto the original function to create a continuous “observed” sequence.

We then coded the sequence as a time series and conducted a time-series analysis in R (version 4.1.2) using the base function `decompose()`. This function separated the waveform into its constituent components: trend, seasonality, and random noise. Trend represented week-long patterns in behavior, seasonality were daily patterns, and random noise was the remaining discrepancy between these two combined effects and the observed sequence.

## Results

### Kruskal-Wallis H

Distributions of fitness metrics generally followed non-gaussian distributions (See Appendix I), and so this required analysis using non-parametric methods. Treatment had no statistical effect on any tested fitness metric (Kruskal-Wallis *H* Test) at an alpha level of 0.05 (Table 2). Therefore, conducting Dunn’s test with Bonferroni Correction as a post-hoc test was unnecessary.

**Table 2.** Kruskal-Wallis *H* Test of fitness vs. treatment level (leaf area).

| **Table 2.** Kruskal-Wallis *H* Test of fitness vs. treatment level (leaf area). |  |  |  |  |
| --- | --- | --- | --- | --- |
| Metric | Chi-squared | df | p-value | Significance |
| Mating Events | 0.51983 | 3 | 0.9145 | n.s. |
| Oviposition Events | 1.9635 | 3 | 0.58 | n.s. |
| Clutches | 0.41339 | 3 | 0.9375 | n.s. |
| Clutch Weight | 0.13509 | 3 | 0.9873 | n.s. |
| Egg Weight | 0.37939 | 3 | 0.9445 | n.s. |
| Percent Hatch | 0.62169 | 3 | 0.8914 | n.s. |

‘n.s.’ indicates $p > 0.05$ .

### Predictive Model and Extrapolation

Because there was no effect of treatment, either *negative* or *positive,* we reasoned that increasing surface area available for flies could be a method to increase number of flies relative to cage volume in a strict sense. Therefore, we summed total perching per cage and fit a linear model with Gaussian distribution to the data to both interpolate and extrapolate how many flies might be added to cages while maintaining the same level of perching behavior (Figure 1).

**Figure 1.**
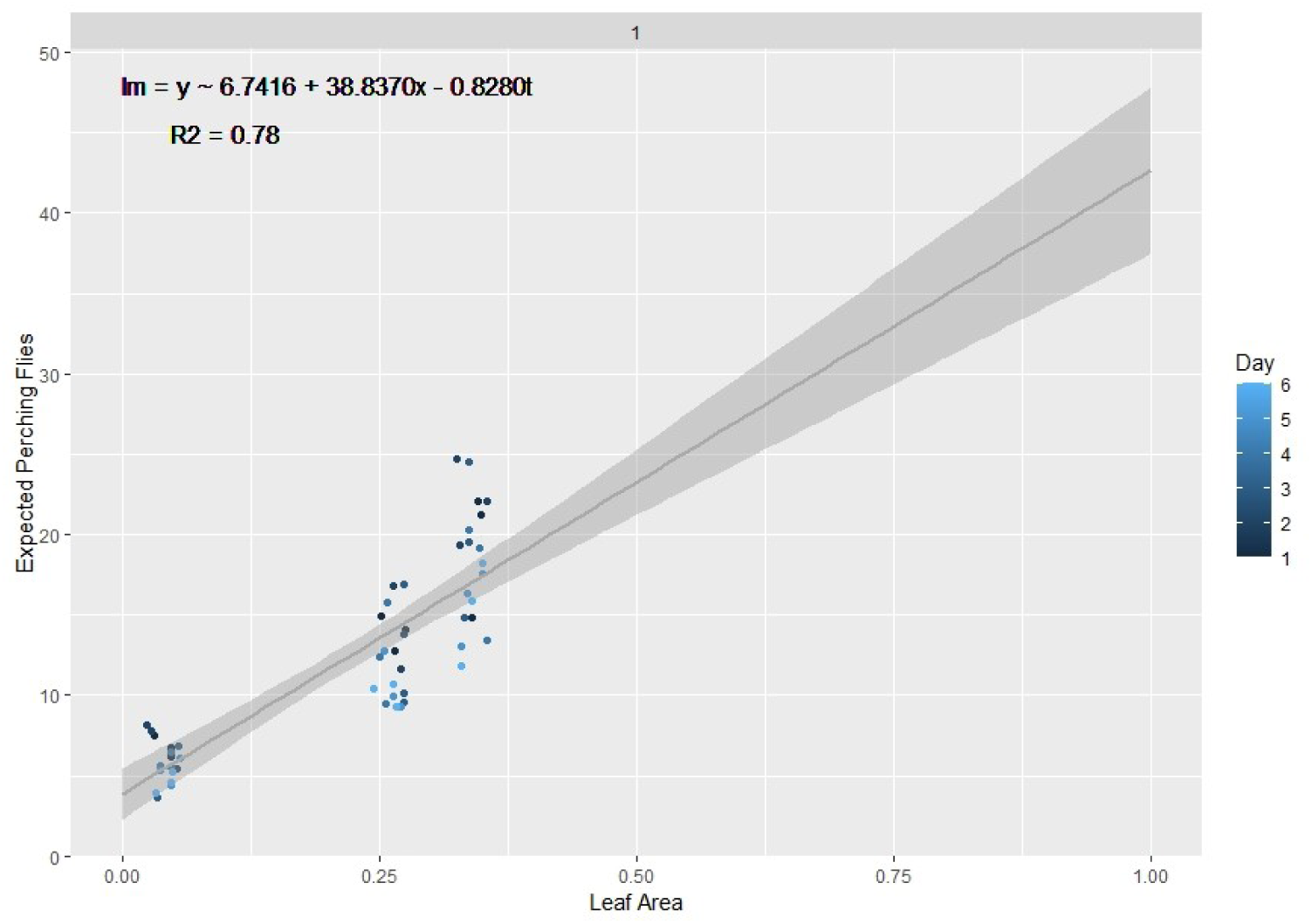
Predictive model relating increases in leaf surface area to percent increase in number of perching black soldier flies. Artificial plants were provided to 0.93 m^3^ cages up to a maximum value of 0.33 m^2^ adaxial surface area. Beyond this, values are extrapolated. Data was also standardized from populations of 90 males and 90 females to 100 of each. The grey area indicates ±95% CI.

The equation for the model can be given as:

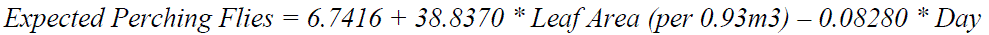

We selected the model parameters (Table 3) by first increasing *R^2^* value (to 0.78) of a linear model and then minimizing Akaike’s Information Criterion (AIC) for the equivalent general linear model (Gaussian family with identity link). We achieved this outcome by, (a) excluding totals from Trial 2, (b) including perching totals from both sexes, and (c) including day as a covariate. We excluded non-significant terms to keep the model parsimonious. We also normalized (i.e., 100 flies per 0.93 m^3^ cage) to align with previous studies (Jones and Tomberlin 2021; Laursen et al. 2022; Dickerson et al. 2024). We then interpolated the results up to a surface area of 1.00 m^2^ (Table 4).

**Table 3.**
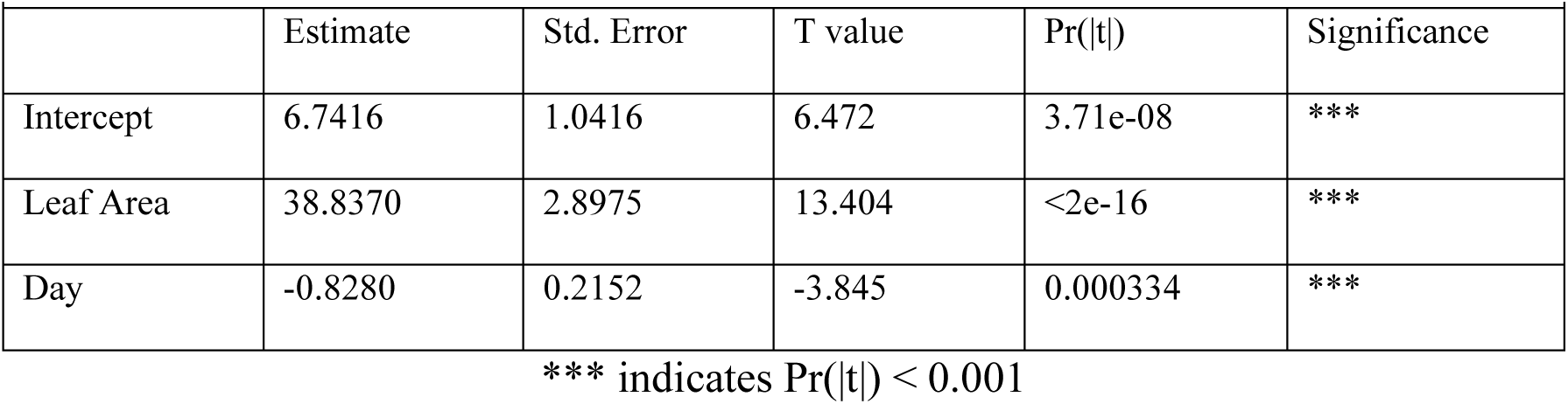
Summary table of linear model. *** indicates Pr(|t|) < 0.001

‘n.s.’ indicates $p > 0.05$ .
| **Table 3.** Summary table of linear model. *** indicates $\Pr(\vert t\vert ) < 0.001$ | | | | | |
| --- | --- | --- | --- | --- | --- |
| | Estimate | Std. Error | T value | $\Pr(\vert t\vert )$ | Significance |
| Intercept | 6.7416 | 1.0416 | 6.472 | 3.71e-08 | *** |
| Leaf Area | 38.8370 | 2.8975 | 13.404 | <2e-16 | *** |
| Day | -0.8280 | 0.2152 | -3.845 | 0.000334 | *** |

\*\*\* indicates $\Pr(|t|) < 0.001$

**Table 4.** Values from predictive model that related increases in added surface area to perching observations. Increases in additional surface area added to 0.93m^3^ breeding cages provided by artificial plants beyond those tested by the experiment are related to increases in number of black soldier flies for a week-long breeding cycle.

|  |  |  |  |  |  |  |  |
| --- | --- | --- | --- | --- | --- | --- | --- |
| Added Surface Area (m<sup>2</sup>/m<sup>3</sup>) | Day 0 | Day 1 | Day 2 | Day 3 | Day 4 | Day 5 | Day 6 |
| 0.25 | 16.45085 | 15.62285 | 14.79485 | 13.96685 | 13.13885 | 12.31085 | 11.48285 |
| 0.50 | 26.1601 | 25.3321 | 24.5041 | 23.6761 | 22.8481 | 22.0201 | 21.1921 |
| 0.75 | 35.86935 | 35.04135 | 34.21335 | 33.38535 | 32.55735 | 31.72935 | 30.90135 |
| 1.00 | 45.5786 | 44.7506 | 43.9226 | 43.0946 | 42.2666 | 41.4386 | 40.6106 |

**Table 5.** Sum of observed female and male perching events per day across n = 3 replicate cages, for each trial. *Indicates in increase in total perching compared to the previous day.

| **Table 5.** Sum of observed female and male perching events per day across n = 3 replicate cages, for each trial. * |  |  |
| --- | --- | --- |
| Indicates in increase in total perching compared to the previous day. |  |  |
| Row Labels | Female Perching | Male Perching |
| Trial 1 | **4675** | **2400** |
| Day 1 | 935 | 366 |
| Day 2 | 961 * | 440 * |
| Day 3 | 779 | 431 |
| Day 4 | 771 | 396 |
| Day 5 | 634 | 389 |
| Day 6 | 595 | 378 |
| Trial 2 | **1124** | **1673** |
| Day 1 | 366 | 276 |
| Day 2 | 285 | 338 * |
| Day 3 | 176 | 224 |
| Day 4 | 184 * | 346 |
| Day 5 | 88 | 365 * |
| Day 6 | 25 | 124 |
| Grand Total | **5799** | **4073** |

### Perch Disparity

While sex disparity was in a strict sense female-biased (Supplementary info), this phenomenon was not static and instead changed over time. On average, the difference between the mean number of females perching on plants and mean number of males perching on plants (defined as ‘perch disparity’, see: Methods) decreased in an absolute sense both over successive days (Figure 2) and hours (Figure 3), towards parity becoming more male biased. For each day of Trial 1, mean (n = 3) perch disparity became more male-biased by 5.78. From 0700 h to 1800 h disparity also shifted towards an additional 2.29 males per cage, per hour on average (n = 3). In Trial 2, sex disparity was initially female biased, but quickly shifted towards male bias on day 2 and later (Figure 5).

**Figure 2.**
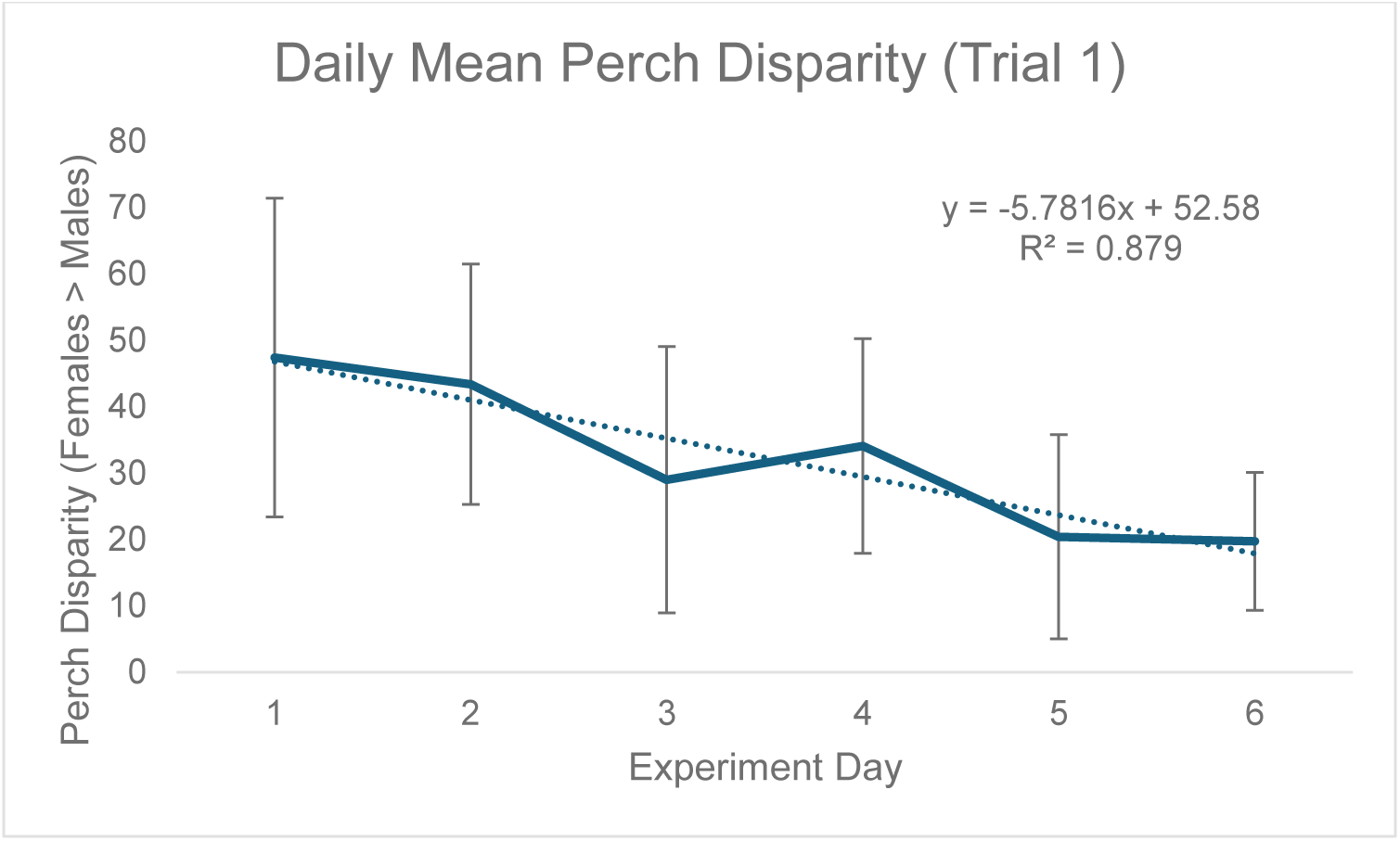
Black soldier fly mean daily perch disparity over time. Perch disparity was calculated as the sum of females perching minus the sum of males perching. Error bars indicate ± SD. Best-fit line is a linear regression. Experimental unit was a 0.93 m^3^ mating cage (n=4 treatments, n = 3 replicates) held within an indoor rearing environment in Texas USA. Each cage had an initial population of 90 males and 90 females.

**Figure 3.**
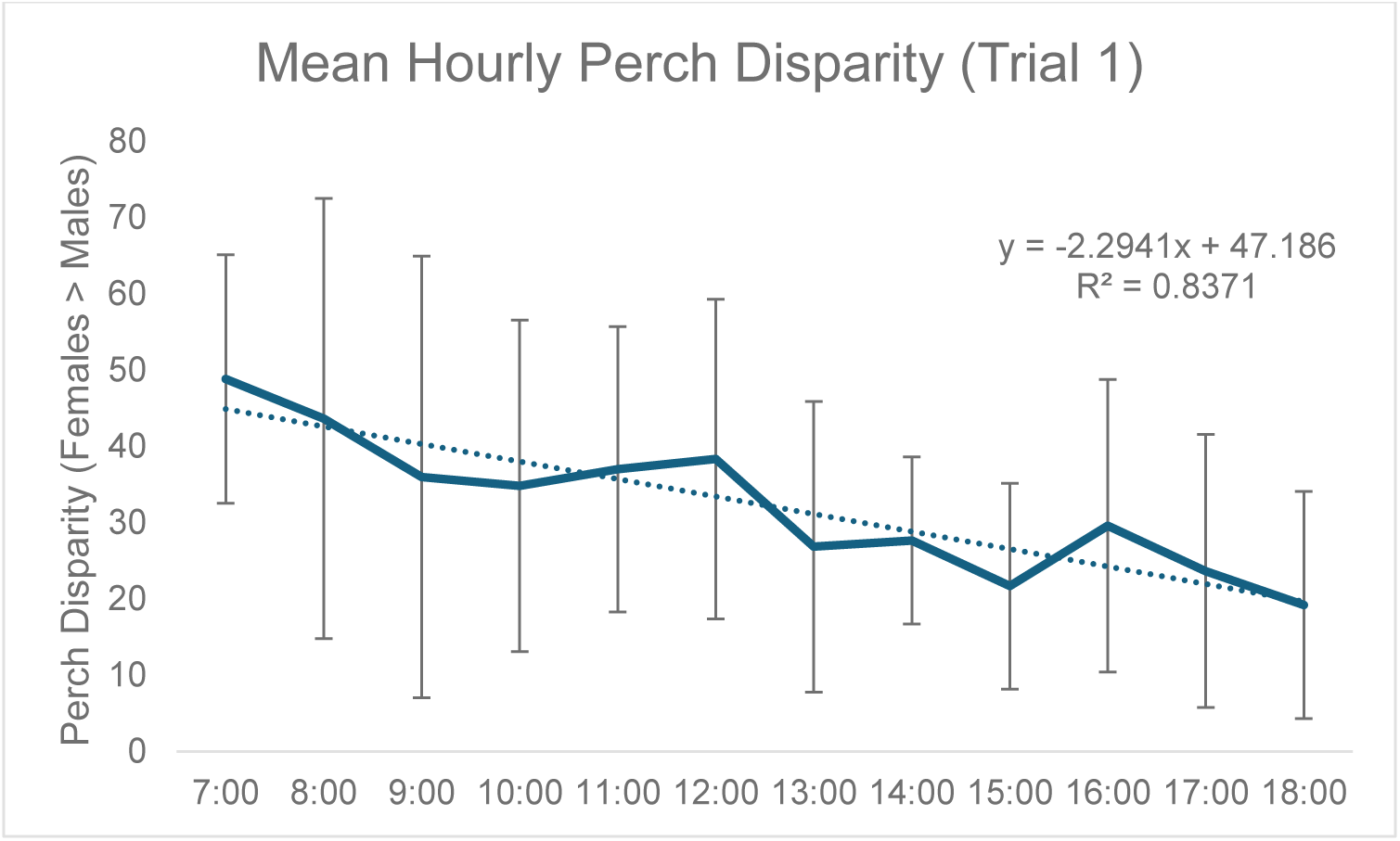
Black soldier fly mean hourly perch disparity over time. Perch disparity was calculated as the sum of females perching minus the sum of males perching. Error bars indicate ± SD. Best-fit line is a linear regression. Experimental unit was a 0.93 m^3^ mating cage (n=4 treatments, n = 3 replicates) held within an indoor rearing environment in Texas USA. Each cage had an initial population of 90 males and 90 females.

Because counts of perching flies were predicated on flies being there to be observed, results were likely impacted by density-effects (e.g., mortality), which were in turn affected by environmental conditions. Although there was an obvious trial effect which caused perch disparity to be much more female biased in trial 1 than in trial 2, both treatment and temporal patterns were consistent between trials (Figure 5). Hence, to approximate a scaling factor between trial 1 and trial 2, we fit a logarithmic regression to both sets of data and compared the intercepts.

Subtracting the two intercepts yielded indicated that an equivalent cage would have been expected to have 57.90 more perching females than males per day in trial 1, as compared to trial 2 (Figure 5). We then conducted time-series analysis on Trial 1-data only (Figure 4) to be more robust to random effects (see supplementary info). From this, results can be generalized to Trial 2 by incorporating the scaling factor of 57.90.

**Figure 4.**
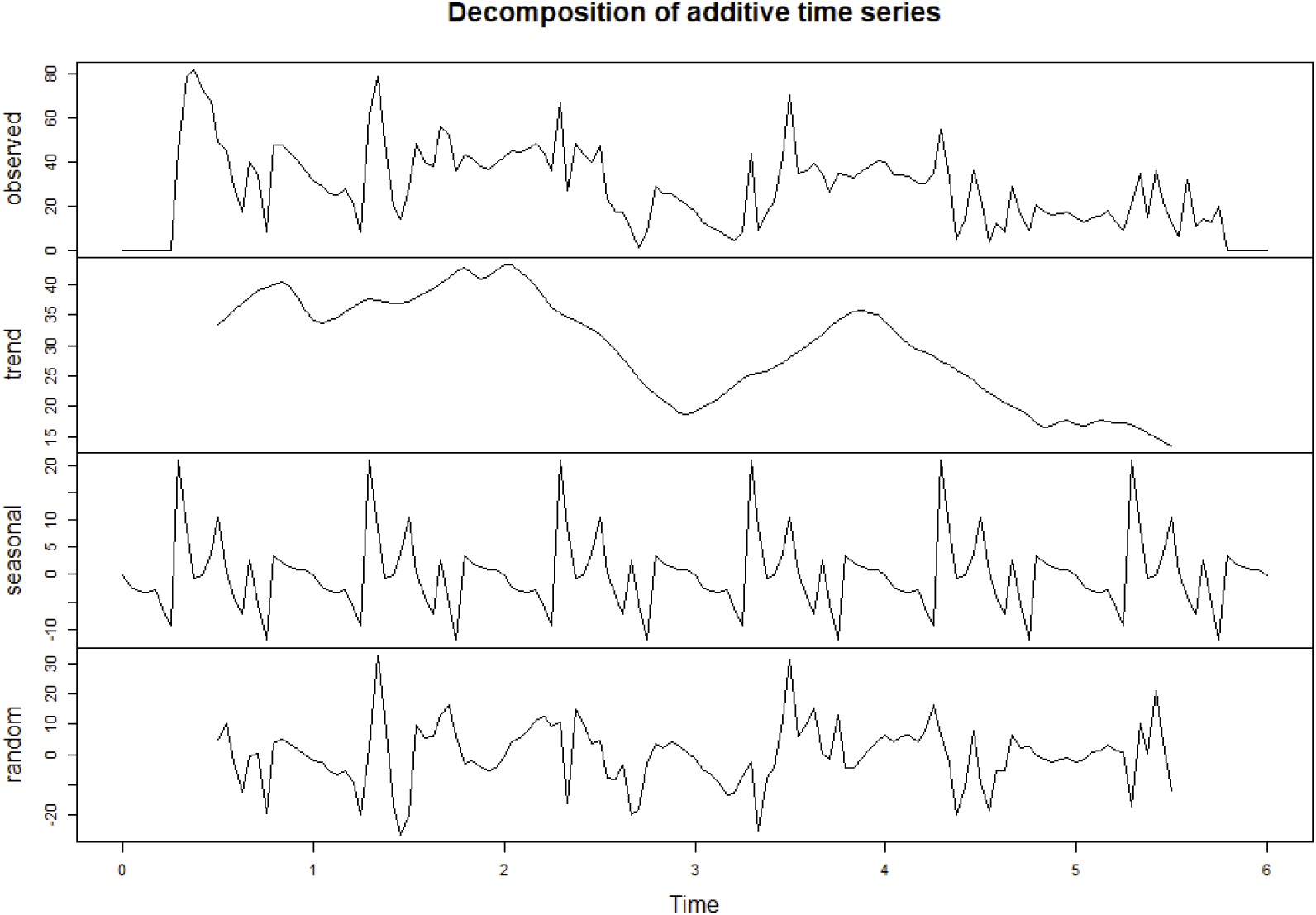
Deconstructed time series analysis of perch disparity. Perch disparity was calculated as the sum of females perching minus the sum of males perching. Data was collected in 0.93 m^3^ mating cages (n = 4 treatments, n = 3 replicates) held in an indoor rearing environment from 0700 h to 1800 h for 6 days. Each cage had an initial population of 90 males and 90 females. Gaps in observed data were filled by taking a 13-hour moving average. Observed data is the sum of ‘trend’ + ‘seasonal’ + ‘random’ data. Respectively, these represent the following: (a) declining perch disparity with each day and a secondary post-mating peak; (b) cyclical pattern of sex bias to peak in the early hours and decline throughout the day; and (c) random effects not otherwise explained.

**Figure 5.**
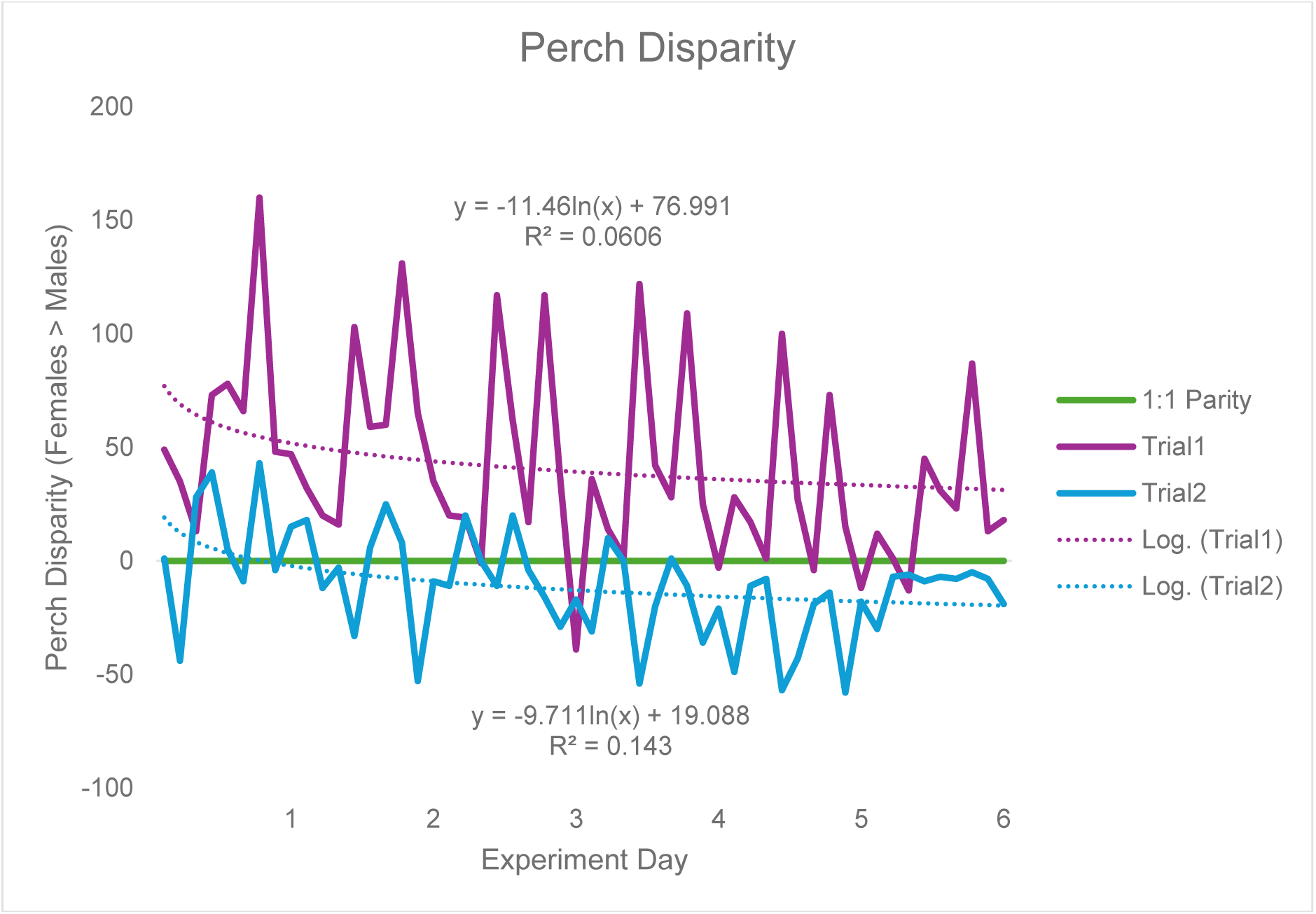
Black soldier perch disparity with respect to trial. Perch disparity was calculated as the sum of females perching minus the sum of males perching, with an x intercept representing parity. Data was collected in 0.93 m^3^ mating cages (n = 4 treatments, n = 3 replicates) held in an indoor rearing environment from 0700 h to 1800 h for 6 days. Each cage had an initial population of 90 males and 90 females. Gaps in observation were not filled in using a moving average. The best fit lines are logarithmic regressions.

### Time Series

Time series analysis (Figure 4) showed that this declination is only temporary, and that female bias recovers by the beginning of observation period each day. The weekly trend has relatively little change in perch disparity during the first three observed days, followed by a large dip towards parity on the third day, a large resurgence in female-bias by day four, then a quick decline in perch disparity that finally reaches parity for the remaining days of the experiment.

## Discussion

By introducing artificial plants of different sizes into breeding cages, this experiment investigated how modifying available surface area to volume ratio can affect fertile egg production of the adult black soldier fly. Because we differentiated females from males by marking their thoraxes with acrylic marker paint (Jones and Tomberlin 2020), we documented sex-specific perching behavior for the first time in captivity. We hypothesized that a medium amount of additional surface area relative to the cage volume would increase fertile egg production, but too much would have a negative effect. In addition, we also thought that males would primarily be those interested in defending territories immediately around perches (Tomberlin and Sheppard 2001). However, treatment had no statistical effect on any recorded fitness metric, either positively or negatively. This suggests that by increasing surface area, additional flies can also be added (Meneguz et al. 2023) without negative effect to improve egg production per cage. Hence, we developed a linear model that could extrapolate the potential additional number of flies that could be accommodated by increasing surface area within cages.

Surprisingly, females engaged in perching much more often than males, though this pattern was dynamic and shifted towards parity throughout time (week/day). In addition, a trial-effect revealed that, in a strict sense, perch disparity (the difference between the number of females and males perching on plants) was density-dependent and strongly affected by environmental conditions. Such conditions were thought to likely have increased morbidity disproportionately in females. This report is especially relevant for producers that are interested in promoting natural behaviors via an insect welfare-minded approach (Boppré and Vane-Wright 2019; Barrett et al. 2022) because presently it is generally unknown which individuals engage in mating versus resting behaviors, at which times during the reproductive cycle, nor how these behaviors might be influenced by the physical structure of the artificial rearing environment.

### Spatial Complexity

Despite our initial hypotheses, this experiment does not provide strong evidence for a link between habitat structural complexity and fitness (McCoy and Bell 1991). We concluded this because mating events, oviposition events, number of clutches, mean clutch weight, harvested egg weight, and hatch percentage (all measured per cage/per day) did not differ significantly across treatments. This finding was contrary to what has been observed in some other flies, particularly mosquitoes like *Aedes agypti* (Linnaeus *in* Hasselquist, 1762) (Diabaté et al. 2011) and *Anopheles gambiae* Giles 1902 (Diptera: Culicidae) (Diabaté et al. 2011). For these species, males form lek-like aerial swarms (Choe and Crespi 1997) near specific physical structures called swarm markers (Diabaté et al. 2011), such that the physical environment can influence variation in mating success. We discuss several potential reasons for this:

First, we designed treatments to crudely approximate the fractal geometry of biological structures and ecological habitats (Williamson and Lawton 1991), rather than test the multi-dimensionality of true spatial complexity. Fractal geometry can be seen in the organization of plant circulatory systems (Weibel 1991), since limbs and roots can continually be sub-divided into self-similar parts irrespective of scale. However, this is only one dimension of spatial complexity which can technically vary based on three separate axes (Williamson and Lawton 1991): (a) heterogeneity, or variation per relative abundance, (b) complexity, or variation per absolute abundance; and (c) scale resolution (Williamson and Lawton 1991). Hence, some dimensions of habitat spatial complexity were not represented in our design. Specifically, our set-up featured plant leaves that were identical copies of one another, and each treatment level was merely a doubling of scale from the previous without change in spatial conformation. Research on perching by Neotropical dung beetles (Coleoptera: Scarabaeidae) suggests that leaf preference varies by insect species and is correlated with individual size, with larger individuals perching on higher and larger leaves (Noriega and Vulinec 2021). Because we found no functional response in terms of fitness to simply increasing the amount of available surface area, this suggests that black soldier flies may instead be more responsive to either an untested structure, or increased range of structures possibly owing to their Neotropical origin (Kaya et al. 2021)where plant structures are hyper-diverse (Barlow et al. 2018).

Second, the level of surface additional surface area relative to the cages (84 × 84 × 133 cm, L × W × H) may simply not have been enough to produce a response to begin with, either positively or negatively. This is because the added surface area for all treatments was less than five percent the total surface area of an empty cage despite occupying a substantial volume. Such is supported by an industrial-scale study that tested additional surface area of 3.33, 3.99, 5.11, and 7.17 m2 per 1.00 m3 cages at densities of 16,000 flies (Grosso et al. 2024). The researchers found that the 3.99 m2/m3 treatment lead to an increased yield of 18.61±4.42 g of eggs compared to 9.23±3.14, 15.00±3.10, and 11.30±3.62 g of eggs for the 3.33, 5.11, and 7.17 m2/m3 treatments; respectively (Grosso et al. 2024). This meant that, for their system, an increase in 0.6 m2/m3 corresponded to a 2.01-fold increase in egg production.

Hilltopping butterflies (Lepidoptera) reportedly avoid areas that are hyper-dense with males and lay few eggs in areas that are spatially bare (Scott 1974). Thus, increasing spatial complexity beyond the range that was tested in this study could promote reproduction by reducing perceived competition and harassment of mated females (Scott 1974; Briceño et al. 1999). While it was hypothesized that too much additional surface would limit reproduction, the potential for artificial plants to be obstacles (visually or spatially) at the scale tested was likely too low.

As fliers, black soldier flies they can easily reposition themselves easily. The amount of additional surface area needed to create a visual or physical barrier when provided by the topology of artificial plants (as opposed to a single, continuous surface—e.g., a wall) is likely much larger for flying insects than for insects that locomote on the ground. Consider that for other farmed insects like crickets (Orthoptera: Ensifera), e.g., *Ruspolia differens* (Serville, 1838) (Orthoptera: Tettigoniidae) (Egonyu et al. 2021), an egg board is added to reduce crowding and provide a visual obstruction to prevent cannibalism (Egonyu et al. 2021), but these insects primarily locomote by crawling or jumping over short distances, and so are not as vagile as black soldier flies.

Third, the choice to use artificial plants over real plants may have removed several elements from the system that are potentially critical for connecting spatial complexity with fitness. These include green-leaf (Metcalf and Kogan 1987) and floral (Knudsen and Gershenzon 2020) volatiles (which are widely known to attract and influence insect behavior), the presence of bacteria (e.g., which the sexually mature male and immature females of flies of the genus *Dacus* (Diptera: Tephritidae) are known to consume (Drew 1987)), as well as different color and thermal properties. While few reports describe plant-insect interactions between adult black soldier flies (e.g., defending territories around Kudzu, *Pueraria montana* (Loer) (Merr.) (Tomberlin and Sheppard 2001), the numerous instances of community science members posting images of black soldier flies on (unidentified) plants (e.g., iNaturalist, BugGuide, LinkedIn, etc.) point to the obviousness and plethora of the interactions that must exist in nature, but at present stand to be documented.

Critically, while the use of real plants theoretically could buffer against environmental stressors by providing shade and reducing critical temperature thresholds—their use, unfortunately, adds an additional layer of complexity, and perhaps unwarranted risk (Lubna et al. 2022), since one of the key issues of growing plants indoors is the threat of pests and pathogens invading without natural enemies to combat them in a relict environment. Our choice to use artificial plants was necessarily born out of the desire to make the experiment replicable and the habitat easier to clean, and mirrors the present Zeitgeist that the transmission of zoonotic pathogens will for the foreseeable future continue to be an issue that affects many organisms living in confined facilities (e.g., confined animal facility operations (Guo et al. 2022)).

### Predictive Model

Because no negative effect of including additional surface area was determined, we fitted a linear model to the data to relate the number of perching flies to increases in leaf area. Extrapolation of model results yielded that, for example increasing surface area by 1.00 m^2^ / 0.93 m^3^ is expected to have the effect of providing enough additional perching area for 45.6 flies at 1:1 sex ratio. While we normalized the model for a density of 200 flies (1:1 sex ratio), it was not normalized to a volume of 1.00 m^3^. This was because the observed patterns, or lack thereof, were likely specific to the cage volume tested. Adding the equivalent surface area to a cage with larger volume or different dimensions is likely to yield different results due to sex-specific lekking behaviors (Lemke et al. 2023) and effects of scale. For instance, the 0.93 m^3^ cage used in this experiment is an order of magnitude larger than, for example, the smallest 0.02 m^3^ cages used by some researchers (Nakamura et al. 2016). It is comparable to some cages used by some industry, while being a third of the size of the relatively standard 2.88 m^3^ cages (EVO Conversion Systems, LLC, Bryan, TX, USA) and two orders smaller the some of the larger 59.42 m^3^ cages available to industry (InsectoCycle, Gelderland, ND, USA). The model results additionally suggest that as days progress, the number of perching flies (viz. females) is expected to diminish slightly, which can be explained by flies expending energy, and becoming less active before dying (personal observation). The precise rate and shape of decline predicted by the model is likely specific to the environmental conditions (see next section) and local adaptations of the tested adult population, but qualitatively should be expected to be generally similar across black soldier flies. For this reason, the results of the model need to be validated independently across a variety of scales.

Additionally, because we placed artificial plants on the bottom of cages and leaves were arranged to maximize exposed surface area, the results should also be interpreted conservatively because we can reasonably assume that flies would have otherwise been able to rest on the floor of the cage. For this reason, future designs should be mindful of ways that additional surface area so that flies have the most opportunity to utilize provided perches, especially when cost is a factor. Future research can explore ways to biologically sort flies through the interaction of behavior with microclimate (Salari and De Goede 2024), for instance, if it is shown that flies vertically stratify based on age or condition within cages, with perhaps the largest, youngest, or most-fit flies preferring different perching locations or temperature-humidity ranges. Similar patterns are well known/documented in a variety of animals, including open-cage and free-range chickens, *Gallus domesticus* (Aves: Galliformes) (Briden et al. 2023), where birds self-stratify based on dominance hierarchies (Haeringen and Hemelrijk 2022). Because insect interactions are size-structured (Alcock 1990; Jones and Tomberlin 2021; Addeo et al. 2022), we have reason to suspect the same can occur for the adult black soldier fly. For this reason, future experiments should consider alternative hypotheses in which flies use all cage surfaces equally or segregate into preferred areas based on underlying biology.

### Perching and Perch Disparity

When considering raw counts alone, females are the sex which that primarily occupied perches. This finding seems to contradict previous description by Tomberlin and Sheppard (Tomberlin and Sheppard 2001) and the general belief held by black soldier fly researchers (personal communication) that males are those to occupy and defend perches to maintain leks. Certainly this response may have been an artifact of the mating cage being restrictive compared to the amount of space black soldier flies likely utilize in the wild (Lemke et al. 2023). Previous studies on perching in insects have documented that males utilize perches to investigate similar-sized objects (i.e., conspecifics) to themselves (Scott 1974). When males encounter other males, this results in brief conflict followed by separation (Scott 1974). When males encounter females this generally resulting in mating (Scott 1974). Similar patterns have been observed in black soldier flies (Tomberlin and Sheppard 2001). Casual observations during our experiment revealed that female black soldier flies intermittently join aerial swarms before returning to their perches.

Returning to the same spot has been described as an energy-saving mechanism because flying to a new spot may result in unwanted harassment (Scott 1974). More generally, insect flight is a highly energetically taxing activity (Beenakkers et al. 1984) that trades-off with egg production (Tanaka 1993; Guerra 2011; Xu et al. 2011)—although different fats are generally used for each (triglyceride vs. phospholipid) (Zhao and Zera 2002). Hence reason dictates that perhaps it is females that gain the most from an increase of available surface area. Sit-and-wait strategies like perching theoretically allow females to reduce total energy usage (Noriega et al. 2020), especially when likely a great deal is needed to constantly maintain flight to and from lekking sites (Roff 1977; Lemke et al. 2023). We speculate the ability to temporarily rest on plants might offer females the ability to preserve precious energy stores while gauging the suitability of mates (Scott 1974), whereas for males constant jostling for position amongst rivals may not have selected for this behavior (Emlen and Oring 1977). Therefore, perching may promote lekking behavior by allowing females a greater opportunity for choice within leks (Alcock 1987), though this needs to be verified. Such behavior can be contrasted with that of *Aedes* mosquitoes who, although engaging in aerial swarms, have limited opportunities for precopulatory female choice other than tarsal kicks (i.e., ‘bucking’) (Aldersley and Cator 2019).

When considering patterns over time in sex disparity data, perching behavior aligns with previous conceptualizations. That is, mating intensity generally peaks in early hours (e.g., by 1200 h under artificial light (Zhang et al. 2010) or 1500 h in natural light (Tomberlin and Sheppard 2002)) and also by the third day of experimentation (Jones and Tomberlin 2021). The amount of mating also negatively regressed with time of day (Tomberlin and Sheppard 2002). Although sex disparity is clearly female-biased, the strength of this bias is dependent on time, and wanes over the course of the day, suggesting that: (a) females could be leaving perches to join mating swarms, or (b) males could be leaving mating swarms to join perches. Both are potentially supported because there are several cases when total perching by either sex increased relative to the preceding day or hour (Tables 6, 7).

**Table 6.** Sum of observed male and female perching events per hour across n = 3 replicate cages, for each trial. *Indicates an increase in total perching compared to the previous hour.

|  | Trial 1 |  | Trial 2 |  |
| --- | --- | --- | --- | --- |
|  | **M** | **F** | **M** | **F** |
| 7:00 AM | 208 | 501 | 145 | 128 |
| 8:00 AM | 238 * | 500 | 128 | 134 * |
| 9:00 AM | 274 * | 490 | 127 | 120 |
| 10:00 AM | 249 | 458 | 127 | 110 |
| 11:00 AM | 287 * | 509 * | 163 | 124 * |
| 12:00 PM | 260 | 490 | 139 | 103 |
| 1:00 PM | 211 | 372 | 164 * | 70 |
| 2:00 PM | 192 | 358 | 196 * | 76 * |
| 3:00 PM | 159 | 289 | 203 * | 84 * |
| 4:00 PM | 101 | 249 | 134 | 70 |
| 5:00 PM | 123 * | 265 | 125 | 67 |
| 6:00 PM | 98 | 194 | 145 * | 128 * |

Critically, the presence of a trial effect revealed that instantaneous operational sex ratio (i.e., the number of males versus females at minute points in time) influenced behavioral patterns, which is in turn was affected by morbidity effects caused by suboptimal abiotic conditions (Scott 1974). In our experiment, although all populations started at an equal 1:1 sex ratio, trial 2 had much higher female mortality rate by the end of the experiment. Each trial experienced differential abiotic effects after the initiation of behavioral observations that likely then caused bottom-up effects on the population structure and behavior, potentially because climactic conditions external to the indoor rearing environment caused the central heating systems to activate more frequently, lowering the relative humidity (See: Supplementary Info).

Specifically, during the grow-out phase, a large difference in the temperature and humidity that flies experienced across trials (i.e., 22.01 °C median temp and 72.27% median RH in trial 1, compared to 26.19 °C median temp and 54.09% median RH in trial 2). Because of this, although the flies used in the experiment were, in a strict sense, the same temporal age, they were unlikely to be the same biological age since both temperature (Holmes et al. 2016; Chia et al. 2018; Addeo et al. 2022) and humidity (Holmes et al. 2012; Bekker et al. 2021; Cammack and Tomberlin 2023) affects the rate at which insects develop. This means that the flies in trial 2 were likely in-effect much older relative to those used in trial 1 (based on relative humidity days) since they experienced more heat and drying days. In addition, after being introduced to cages, although starting at the same RH across-trials, the breeding environment in trial 1 increased in relative humidity, while the breeding environment in trial 2 decreased in humidity, compounding the difference by 14.7 RH-days (Supplementary Info).

Past studies have shown differing results regarding longevity with either: (a) females living ∼3.5 less days than males irrespective of rearing temperature, or (b) no difference between male and females (Klüber et al. 2023), potentially due to the ability of poikilotherms to thermoregulate (Addeo et al. 2022). Alternatively another study suggested that, by mating, females increase their longevity relative to males by gaining nutrition via nuptial gifts (Harjoko et al. 2023), this is potentially supported by our experiment, especially if the extension of female longevity is predicated on females mating early in their (biological) lives. If flies were held in holding cages and were delayed mating (Dickerson et al. 2024), perhaps this then explains the increased mortality of females in Trial 2; however, no such effect has been seen in other experiments on black soldier fly aging (Dickerson et al. 2024; Lemke 2024).

This study reveals that behavior is complex and not easily intuited. There is still much to be understood, even in environments that are purportedly environmentally controlled (Li et al. 2023). As the field continues to progress and describe these multi-tiered interactions, it is important that both theory and empiricism continue to be linked and inform one another.

### Lekking?

While a 0.93-m^3^ cage has often been thought to be too crowded to promote lekking behavior (Lemke et al. 2023), the provisioning of our experimental cages with artificial plants provided the opportunity to show that perhaps at least some elements of black soldier fly-lekking behavior is preserved within captivity, especially in larger cages; though this does not rule out the possibility that lekking is being selected against in favor of something akin to scramble-competition polygyny (Emlen and Oring 1977; Dodson 1986).

Identifying criteria (Bradbury et al. 1986) for lekking have been formalized and applied to insects (Alcock 1987; Choe and Crespi 1997) including the black soldier fly (Tomberlin and Sheppard 2001; Birrell 2018), the closely related *Hermetia comstockii* Williston (Diptera: Stratiomyidae) (Alcock 1990), as well as orchid bees (Hymenoptera: Apinae: Euglossini) (Kimsey 1980). The aspects of lekking that seem to be preserved for the black soldier fly in captivity include the spatiotemporal segregation of males and females since males primarily occupied aerial swarms during early hours and females were those that primarily occupied perches. In addition, males engaged in prolonged bouts of mating behavior near lights, as indicated by casual observations. These mating swarms occurred in cages regardless of whether perches or attractant box were present, suggesting that the mating system is unlikely to be akin to resource-defense (Hastings et al. 1994) or female-defense polygyny. Instead, the structure of black soldier fly leks (i.e., their dimensions) appear quite plastic and may contract or grow depending on the available space (i.e., exploded vs. imploded leks) (Höglund and Alatalo 1995).

This study’s results also suggest that any positive welfare effect of increasing surface area will likely be primarily towards females, as when environmental conditions were optimal, they were those that utilized perching structures the most. Future research should aim to devise ethograms (e.g., (Kimsey 1980; Briceño et al. 1999; Masse et al. 2023)) to properly quantify the positive contribution of perches to improving quality-of-life. Although there was not a direct link between fitness and increased surface area quantified (albeit keeping population density constant), the placement of the artificial structures allowed for the re-conceptualization of what can occur within the confines of even a 0.93-m^3^ cage, such that the lek-mating system may still persists within captivity for black soldier flies (for now) despite the pressures of artificial selection (Price 1984). Indeed, lekking may be encouraged through the modification of the artificial habitat (Liedo et al. 2007). It is possible with a larger space, separate areas can be devised for each sex—one in which males can engage in mating bouts, and another for females to rest, continue their reproductive development, and lay eggs. Certainly, prior work has implicated the presence of old males in swarms to either have no positive, or a net negative contribution to fitness (Dickerson et al. 2024; Lemke 2024), and so devising ways to isolate the non-reproductive from the reproductive flies within cages will like promote applied aims of getting consistent egg-production.

