## Supplementary Info for "Sex-specific perching: Monitoring of artificial plants reveals dynamic female-biased perching behavior in the black soldier fly, *Hermetia illucens* (Diptera: Stratiomyidae)"

Supplementary Info I

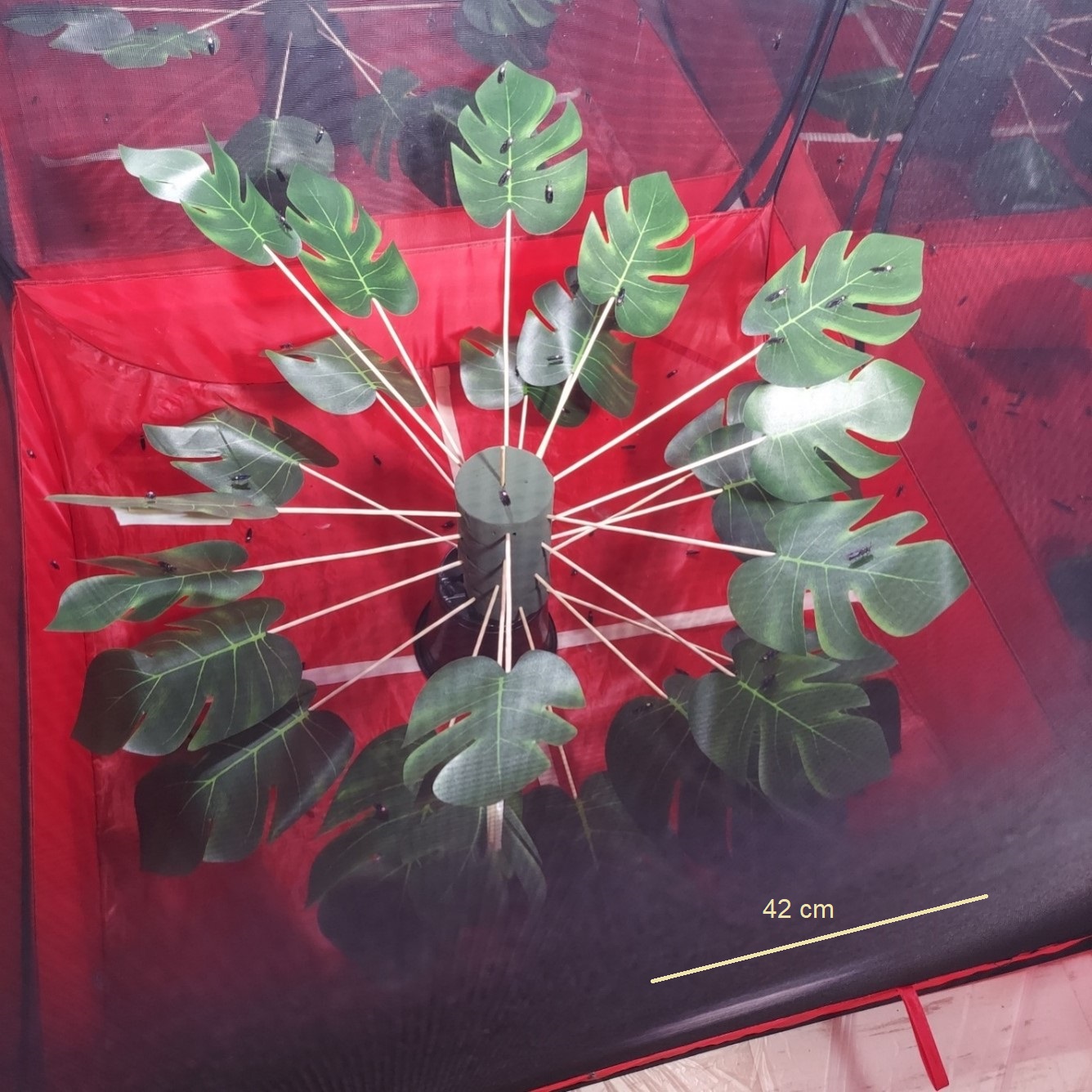

Figure 10. Black soldier flies perching on artificial *Monstera deliciosa* plant in HIGH treatment. The experimental unit was a 0.93m^3^ breeding cage housed within the FLIES Facility, Texas A&M University, USA. Each cage had an initial population of 90 males and 90 females. Females are indicated by pink dot of acrylic marker paint on thorax, while males have no marking.

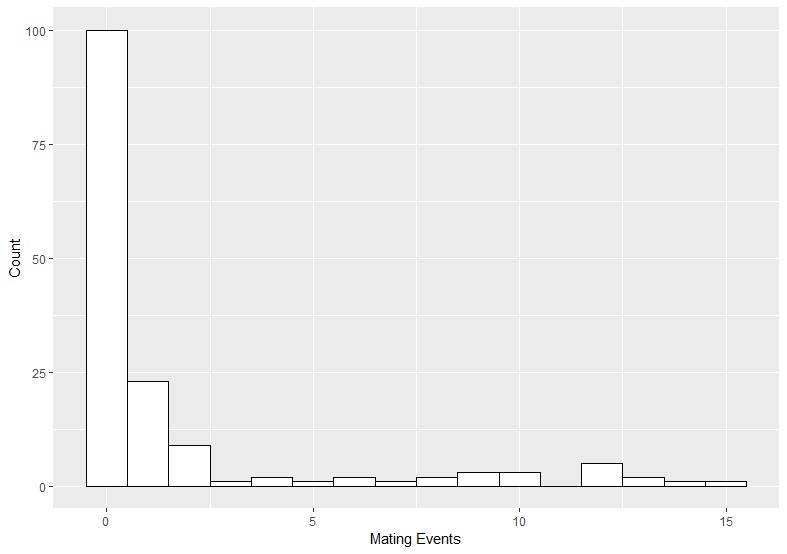

Figure 11. Histogram of observed mating events. Counts were combined across two trials, conducted indoors at the FLIES facility. Observations of black solider flies were recorded hourly from 0700-1600 h, for 6 d, in populations of 90 males and 90 females in 0.93 m3 cages.

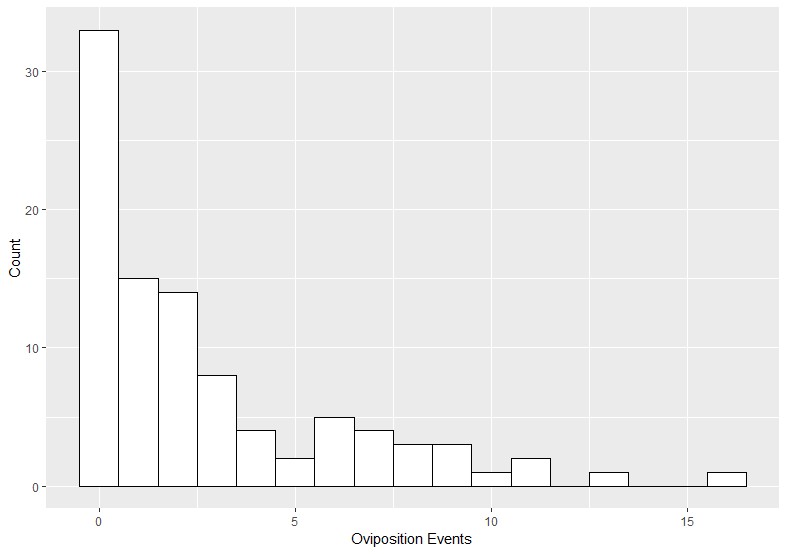

Figure 12. Histogram of observed oviposition events. Counts were combined across two trials, conducted indoors at the FLIES facility. Observations of black solider flies were recorded hourly from 0700-1600 h, for 3 d starting after the introduction of attractant boxes on the third day of experimentation, in populations of 90 males and 90 females in 0.93 m^3^ cages.

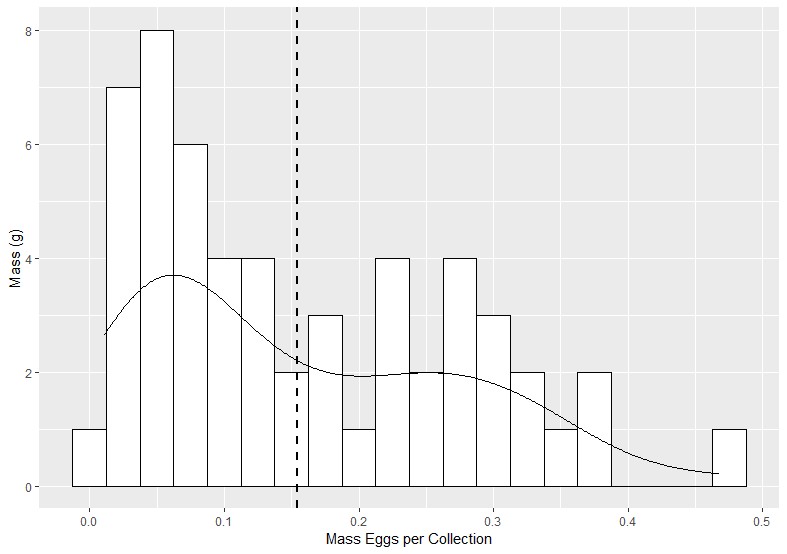

Figure 13. Mass of eggs collected per cage, per day. Eggs were harvested from cardboard egg traps daily, each day of experimentation after the third day. All eggs laid in a single trap constituted a cage-day replicate. The vertical dashed line in the figure represents the mean mass of eggs collected, agnostic of replicate. A smoothing function is overlaid over the histogram, and suggests a bimodal distribution, with local maxima occurring to either side of the mean.

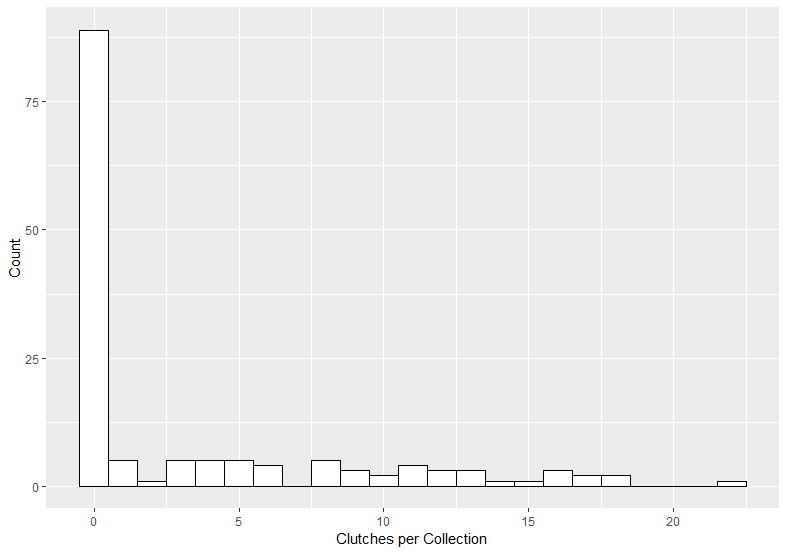

Figure 14. Count of clutches laid by black soldier flies within cardboard egg traps. Clutches were identified as being distinct and physically separable masses. Eggs were harvested after the third day, from each cage, daily.

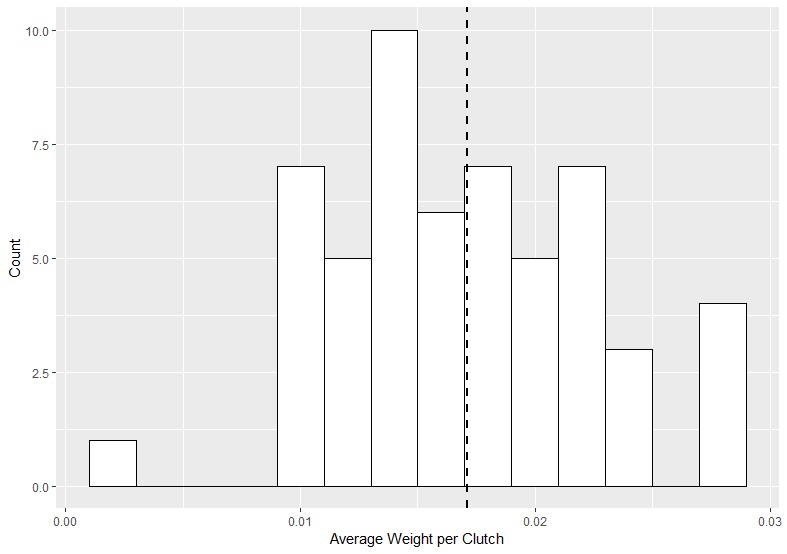

Figure 15. Mean ‘clutch’ weight of eggs collected. Clutches were identified as egg masses which were physically separable from one another within cardboard egg traps. Mean clutch weight was determined by taking the count of clutches and dividing that number into the weight of eggs collected, per cage, per day from 0.93 m^3^ breeding cages within initial populations of 90 males and 90 females. The vertical dashed line indicates the global mean.

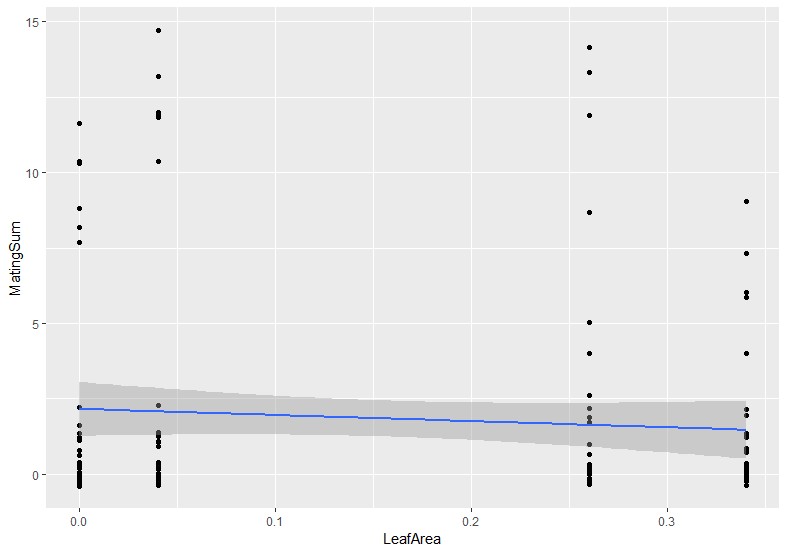

Figure 16. Linear regression of summed Mating Events in response to increasing adaxial leaf surface area. The grey area represents ±95% CI. Artificial *Monstera deliciosa* leaves were provided to breeding populations of 90 male and 90 female black soldier flies in 0.93 m^3^ cages housed indoors within the FLIES Facility.

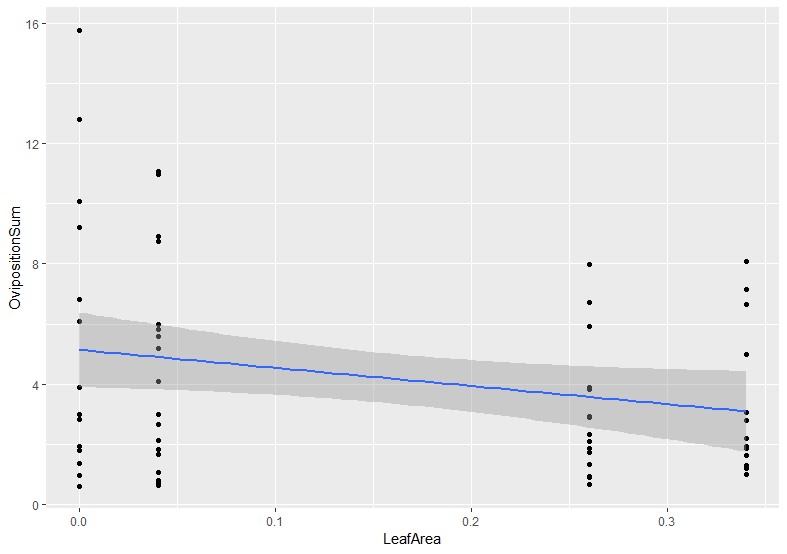

Figure 17. Linear regression of summed observed Oviposition Events in response to increasing adaxial leaf surface area. The grey area represents ±95% CI. Artificial *Monstera deliciosa* leaves were provided to breeding populations of 90 male and 90 female black soldier flies in 0.93 m^3^ cages housed indoors within the FLIES Facility.

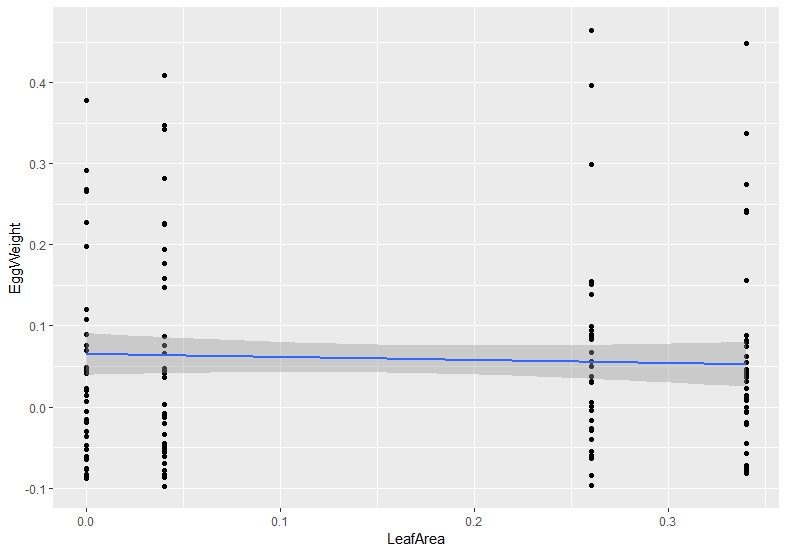

Figure 18. Regression of summed Egg Weight in response to increasing adaxial leaf surface area. The grey area represents ±95% CI. Artificial *Monstera deliciosa* leaves were provided to breeding populations of 90 male and 90 female black soldier flies in 0.93 m^3^ cages, housed indoors within the FLIES Facility.

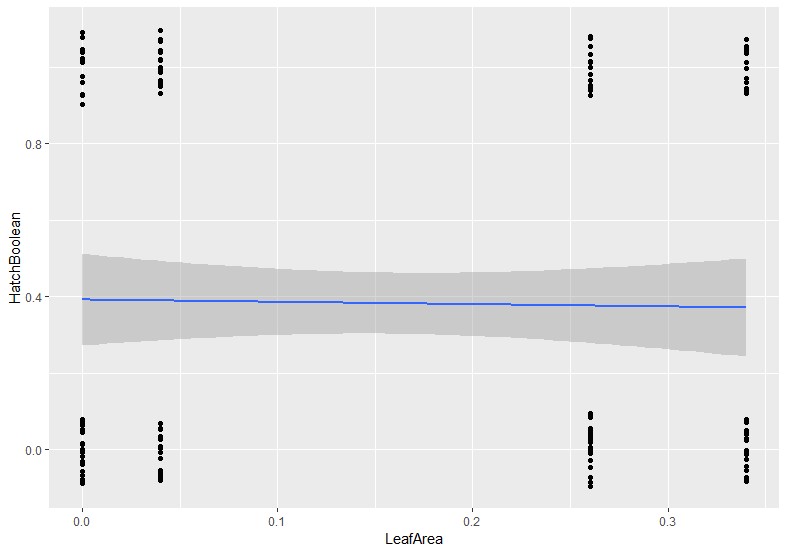

Figure 19. Linear regression of Hatch Rate as Boolean response to increasing adaxial leaf surface area. The grey area represents ±95% CI. Artificial *Monstera deliciosa* leaves were provided to breeding populations of 90 male and 90 female black soldier flies in 0.93 m^3^ cages.

Supplemantary Info II

Analysis of Differential Abiotic Effects

Because there was substantial mortality during the second trial, but not the first trial, I conducted several additional analyses to determine what was likely the root cause of this morbidity. The results generally indicate that across trials flies experienced similar temperatures, but drastically different humidity, and both were kept in conditions that were likely suboptimal.

Summarized Methods

*Data captured from HOBO Logger placed in Room 9
*Plotted using Excel
*Best-fit regressions selected based on R2 values (increasing beyond a 4^th^-order polynomial did not significantly improve R2 for one of the models but did for the other).
*The area under the curve was found using Wolfram Alpha (i.e., difference between two integrals) which represented the accumulated degree/humidity days.

Results

Temperature x Trial

Temperature

Degrees C

| Experiment Day | Trial 1 | Trial 2 |
| --- | --- | --- |
| 0 | 34.119 | 30.4371 |
| 0 | 32.8053 | 30.8988 |
| 0 | 33.3115 | 31.2322 |
| 0 | 34.0484 | 30.8529 |
| 0 | 33.024 | 31.5859 |
| 0 | 33.0467 | 30.9163 |
| 1 | 34.2357 | 30.4671 |
| 1 | 34.2061 | 31.0432 |
| 1 | 33.8206 | 31.2752 |
| 1 | 32.8328 | 31.7856 |
| 1 | 33.024 | 31.2688 |
| 1 | 34.2653 | 31.229 |
| 1 | 33.9828 | 31.0781 |
| 1 | 32.8458 | 31.0463 |
| 1 | 32.0644 | 31.1702 |
| 1 | 33.6897 | 31.5333 |
| 1 | 34.0468 | 30.8846 |
| 1 | 32.5534 | 30.2763 |
| 1 | 32.4373 | 30.4087 |
| 1 | 32.6663 | 30.445 |
| 1 | 29.2243 | 30.4402 |
| 1 | 32.2908 | 30.0276 |
| 1 | 31.8112 | 31.1511 |
| 1 | 30.6345 | 31.1289 |
| 1 | 31.3086 | 30.6187 |
| 1 | 31.9185 | 30.3235 |
| 1 | 31.5492 | 31.0797 |
| 1 | 32.2362 | 30.1361 |
| 1 | 30.693 | 31.0765 |
| 1 | 32.3101 | 31.2322 |
| 2 | 31.7856 | 30.2526 |
| 2 | 31.1845 | 30.8497 |
| 2 | 32.1896 | 30.8513 |
| 2 | 32.4067 | 31.2799 |
| 2 | 33.1522 | 31.0225 |
| 2 | 31.5556 | 30.9448 |
| 2 | 31.6993 | 30.948 |
| 2 | 31.7025 | 30.9131 |
| 2 | 31.7041 | 31.3834 |
| 2 | 31.9762 | 30.5049 |
| 2 | 31.4408 | 31.3755 |
| 2 | 32.9737 | 31.3038 |
| 2 | 32.655 | 30.0685 |
| 2 | 32.0756 | 30.3819 |
| 2 | 31.5428 | 30.7262 |
| 2 | 32.8426 | 28.6454 |
| 2 | 32.7568 | 29.318 |
| 2 | 30.0245 | 29.1978 |
| 2 | 32.0997 | 29.0575 |
| 2 | 31.094 | 29.137 |
| 2 | 32.3391 | 29.5791 |
| 2 | 30.1645 | 29.19 |
| 2 | 31.5524 | 30.1597 |
| 2 | 31.1575 | 29.5572 |
| 3 | 31.3436 | 30.0339 |
| 3 | 31.1813 | 29.3617 |
| 3 | 31.1432 | 29.7247 |
| 3 | 31.8272 | 30.4039 |
| 3 | 32.005 | 29.7498 |
| 3 | 32.476 | 29.9491 |
| 3 | 32.6195 | 30.5934 |
| 3 | 31.1241 | 30.1393 |
| 3 | 31.8432 | 29.6605 |
| 3 | 32.2619 | 30.0921 |
| 3 | 30.6203 | 30.1 |
| 3 | 31.9698 | 30.3472 |
| 3 | 30.7515 | 29.1807 |
| 3 | 31.0447 | 30.2448 |
| 3 | 29.72 | 29.1931 |
| 3 | 31.5317 | 29.9036 |
| 3 | 29.626 | 29.7232 |
| 3 | 31.8496 | 29.7436 |
| 3 | 27.5664 | 29.393 |
| 3 | 30.0056 | 28.8412 |
| 3 | 29.3383 | 29.9507 |
| 3 | 28.3786 | 30.4544 |
| 3 | 28.8458 | 29.3477 |
| 3 | 29.9193 | 30.1613 |
| 4 | 29.4743 | 29.2415 |
| 4 | 28.6749 | 29.9475 |
| 4 | 29.1604 | 30.2983 |
| 4 | 28.9283 | 30.0292 |
| 4 | 28.8878 | 29.2321 |
| 4 | 28.9034 | 29.4899 |
| 4 | 29.2961 | 30.0544 |
| 4 | 29.601 | 30.3093 |
| 4 | 30.0528 | 29.2197 |
| 4 | 30.0685 | 29.9931 |
| 4 | 30.4528 | 29.4602 |
| 4 | 28.8816 | 29.8282 |
| 4 | 30.0968 | 30.111 |
| 4 | 30.218 | 30.0968 |
| 4 | 30.3551 | 28.6143 |
| 4 | 28.0491 | 29.4555 |
| 4 | 28.012 | 29.8737 |
| 4 | 27.608 | 28.9874 |
| 4 | 27.0594 | 29.7796 |
| 4 | 27.1698 | 29.9161 |
| 4 | 27.1683 | 29.8533 |
| 4 | 27.0716 | 29.5368 |
| 4 | 26.8571 | 29.6323 |
| 4 | 26.8418 | 29.7671 |
| 5 | 26.6535 | 29.7639 |
| 5 | 26.6337 | 29.8 |
| 5 | 26.5908 | 30.0764 |
| 5 | 26.6704 | 30.0764 |
| 5 | 26.6474 | 30.0497 |
| 5 | 26.4136 | 29.789 |
| 5 | 26.7285 | 29.88 |
| 5 | 26.464 | 30.1975 |
| 5 | 26.4701 | 30.2605 |
| 5 | 26.4228 | 30.0559 |
| 5 | 26.1512 | 28.5988 |
| 5 | 26.0552 | 29.4117 |
| 5 | 26.0964 | 30.4055 |
| 5 | 26.1467 | 28.6858 |
| 5 | 26.3846 | 29.7044 |
| 5 | 26.2488 | 30.0135 |
| 5 | 26.1512 | 30.0276 |
| 5 | 30.1897 | 28.4794 |
| 5 | 26.8448 | 29.1121 |
| 5 | 25.9182 | 28.9594 |
| 5 | 25.7463 | 30.155 |
| 5 | 25.5822 | 29.6057 |
| 5 | 25.3849 | 30.2101 |
| 5 | 25.309 | 30.3109 |
| 6 | 25.2196 | 29.9538 |
| 6 | 25.0438 | 29.1994 |
| 6 | 25.0196 | 30.576 |
| 6 | 25.0787 | 30.133 |
| 6 | 25.0287 | 29.4852 |
| 6 | 24.9182 | 29.902 |
| 6 | 25.109 | 30.3598 |
| 6 | 25.3575 | 31.1957 |
| 6 | 25.4122 | 30.155 |
| 6 | 25.5214 | 30.6092 |
| 6 | 28.7743 | 30.8624 |
| 6 | 28.9034 | 30.5365 |
| 6 | 29.1027 | 29.9507 |
| 6 | 29.3992 | 30.4071 |
| 6 | 27.1315 | 29.6997 |
| 6 | 27.9533 | 30.3961 |
| 6 | 28.8085 | 29.9774 |
| 6 | 27.1576 | 29.5009 |
| 6 | 27.6418 | 29.9962 |
| 6 | 28.2192 | 29.9114 |
| 6 | 28.2811 | 29.8879 |
| 6 | 28.2285 | 29.7733 |
| 6 | 27.9734 | 30.5523 |
| 6 | 28.2114 | 30.3157 |

Figure 23. ­­Relative degree day difference experienced by flies during Trial 1 and Trial 2. The most parsimonious polynomial regressions were fit to time series data, and then the area between the curves was calculated by taking the difference of the two integrals using Wolfram Alpha.

Adults in Trial 1 experienced an estimated 10.51 more-degree days than trial two, until midway through day 3 of the experiments, after which, trial two experienced an estimated 6.40 more-degree days than trial one. Although the maximum temperature T1 flies experienced was higher, this coincided with increased performance, particularly during the period of life when mating and oviposition were most likely to occur (i.e., before day 4) – and so it is unlikely that increased temperatures have a large effect on the behavioral patterns seen in T2.

Humidity x Trial

Instead, the major contributing factor to the decline in perching activity, and adult mortality in T2 is lack of humidity (measured by % RH).

Trial 1 experienced ~14.8 more relative humidity days (or up to 82 if day 0 is included), even though at the start of both trials, relative humidity was equal at around 40% RH. Trends show that over time, humidity for Trial 1 increased, while in Trial 2 it decreased.

Relative Humidity

% RH

| Experiment Day | Trial 1 | Trial 2 |
| --- | --- | --- |
| 0 | 38.3438 | 38.1332 |
| 0 | 38.7146 | 37.9471 |
| 0 | 38.913 | 37.5443 |
| 0 | 41.1728 | 36.9232 |
| 0 | 40.3687 | 35.199 |
| 0 | 41.4856 | 34.5612 |
| 1 | 41.5039 | 34.3552 |
| 1 | 40.7532 | 33.165 |
| 1 | 40.1016 | 32.4371 |
| 1 | 40.033 | 31.4301 |
| 1 | 40.2542 | 30.7343 |
| 1 | 39.7079 | 29.9881 |
| 1 | 38.7604 | 30.9082 |
| 1 | 39.4714 | 29.9103 |
| 1 | 38.7939 | 29.8508 |
| 1 | 39.5081 | 30.304 |
| 1 | 38.2217 | 30.2917 |
| 1 | 37.1521 | 30.9235 |
| 1 | 37.0377 | 31.2698 |
| 1 | 36.998 | 31.0684 |
| 1 | 41.1591 | 30.864 |
| 1 | 40.2344 | 34.5398 |
| 1 | 41.7267 | 32.0801 |
| 1 | 40.39 | 33.3527 |
| 1 | 42.9535 | 31.9473 |
| 1 | 40.9271 | 31.2653 |
| 1 | 39.2227 | 31.0425 |
| 1 | 37.1124 | 30.6107 |
| 1 | 36.5433 | 29.7562 |
| 1 | 36.174 | 29.9606 |
| 2 | 35.6384 | 29.7699 |
| 2 | 35.7941 | 29.1519 |
| 2 | 34.8236 | 28.9902 |
| 2 | 34.9411 | 28.9764 |
| 2 | 34.5764 | 28.2623 |
| 2 | 35.3714 | 27.916 |
| 2 | 34.4208 | 27.832 |
| 2 | 40.7059 | 29.66 |
| 2 | 40.4724 | 29.0421 |
| 2 | 38.8977 | 28.8666 |
| 2 | 36.7279 | 29.4678 |
| 2 | 35.1944 | 29.4357 |
| 2 | 41.3025 | 30.896 |
| 2 | 38.8763 | 31.9016 |
| 2 | 37.291 | 35.8582 |
| 2 | 36.6257 | 35.0113 |
| 2 | 35.7147 | 33.0246 |
| 2 | 42.8085 | 34.465 |
| 2 | 39.801 | 35.1471 |
| 2 | 37.5488 | 33.9706 |
| 2 | 35.6934 | 33.2031 |
| 2 | 35.6155 | 33.1253 |
| 2 | 34.8602 | 32.6538 |
| 2 | 33.5785 | 32.3608 |
| 3 | 32.7408 | 31.9427 |
| 3 | 33.4488 | 32.2556 |
| 3 | 33.8104 | 31.9748 |
| 3 | 33.4686 | 31.6879 |
| 3 | 33.638 | 31.4575 |
| 3 | 33.4442 | 31.2012 |
| 3 | 33.1787 | 31.4407 |
| 3 | 39.3753 | 31.9427 |
| 3 | 38.5513 | 32.2281 |
| 3 | 36.7355 | 32.4463 |
| 3 | 36.1694 | 32.579 |
| 3 | 34.8877 | 32.3242 |
| 3 | 33.963 | 32.5607 |
| 3 | 41.3208 | 32.3135 |
| 3 | 38.2431 | 31.4682 |
| 3 | 34.964 | 32.4554 |
| 3 | 35.08 | 31.073 |
| 3 | 40.8798 | 31.5689 |
| 3 | 44.8944 | 32.2037 |
| 3 | 40.1794 | 32.6721 |
| 3 | 38.5147 | 32.2144 |
| 3 | 38.4506 | 32.8003 |
| 3 | 37.5473 | 33.0338 |
| 3 | 36.9339 | 33.5358 |
| 4 | 38.0096 | 34.1187 |
| 4 | 37.9181 | 34.1904 |
| 4 | 37.7731 | 34.52 |
| 4 | 38.0142 | 35.2859 |
| 4 | 38.2858 | 35.3134 |
| 4 | 38.4766 | 35.1379 |
| 4 | 37.9089 | 34.9319 |
| 4 | 44.4153 | 36.9293 |
| 4 | 44.635 | 36.7508 |
| 4 | 41.9968 | 35.4904 |
| 4 | 39.9734 | 36.6089 |
| 4 | 40.1398 | 35.5774 |
| 4 | 44.5557 | 36.351 |
| 4 | 40.8798 | 36.6806 |
| 4 | 39.7079 | 36.8729 |
| 4 | 40.3397 | 36.0825 |
| 4 | 40.1337 | 36.5082 |
| 4 | 46.9315 | 36.2976 |
| 4 | 45.6131 | 36.7691 |
| 4 | 44.0811 | 37.3917 |
| 4 | 42.7841 | 37.3505 |
| 4 | 42.0944 | 37.5793 |
| 4 | 41.7831 | 37.8281 |
| 4 | 42.4881 | 38.5269 |
| 5 | 42.4332 | 38.9404 |
| 5 | 42.4591 | 39.3066 |
| 5 | 42.8986 | 40.0299 |
| 5 | 43.5089 | 40.6113 |
| 5 | 43.9072 | 41.2369 |
| 5 | 43.8873 | 41.0507 |
| 5 | 44.0674 | 41.1743 |
| 5 | 50.4807 | 42.0609 |
| 5 | 49.0112 | 42.0013 |
| 5 | 48.9182 | 40.2679 |
| 5 | 48.7411 | 39.8972 |
| 5 | 47.67 | 39.1602 |
| 5 | 56.9717 | 38.8885 |
| 5 | 53.9169 | 40.7883 |
| 5 | 50.9186 | 38.7695 |
| 5 | 49.2157 | 38.5269 |
| 5 | 47.5784 | 38.6108 |
| 5 | 39.0717 | 38.8123 |
| 5 | 44.4092 | 37.5381 |
| 5 | 45.4208 | 37.3779 |
| 5 | 46.6354 | 36.9507 |
| 5 | 46.7392 | 35.8307 |
| 5 | 46.8124 | 35.6567 |
| 5 | 47.6685 | 34.4666 |
| 6 | 47.1985 | 33.1528 |
| 6 | 48.3414 | 33.0521 |
| 6 | 48.6252 | 32.3639 |
| 6 | 48.6206 | 31.1066 |
| 6 | 48.7411 | 30.4688 |
| 6 | 49.2218 | 29.1107 |
| 6 | 50.2747 | 28.685 |
| 6 | 52.2156 | 28.2211 |
| 6 | 52.5757 | 27.742 |
| 6 | 55.8731 | 27.713 |
| 6 | 43.7408 | 27.6596 |
| 6 | 44.931 | 27.063 |
| 6 | 49.3301 | 27.2278 |
| 6 | 47.995 | 26.6785 |
| 6 | 48.7991 | 27.2202 |
| 6 | 46.3089 | 26.7395 |
| 6 | 43.3151 | 26.4282 |
| 6 | 50.676 | 26.6434 |
| 6 | 47.8897 | 26.326 |
| 6 | 46.4691 | 26.0696 |
| 6 | 45.5139 | 25.6714 |
| 6 | 45.0684 | 25.0763 |
| 6 | 44.3771 | 25.2182 |
| 6 | 44.7159 | 24.8337 |

Figure 24. Relative difference in drying days experienced by flies during Trial 1 and Trial 2. The most parsimonious polynomial regressions were fit to time series data, and then the area between the curves was calculated by taking the difference of the two integrals using Wolfram Alpha.

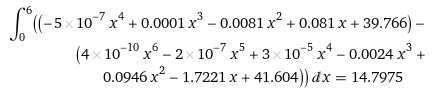

Figure 25. Integral expressing the area (difference) between the two calculated best-fit lines, yielding difference in ‘humidity’ or ‘drying’ days.

If optimal ambient humidity is (conservatively) ~60% (but up to 70%, See: Nakamura et al. 2016), then this means trial 1 and trial 2 experienced 120.5 or 135.9 fewer humidity days than would have been required for peak performance, respectively. And so, with that in mind, the interpretation of the data should reflect that Trial 1 occurred in conditions in which flies tolerated, while Trial 2 occurred in conditions that were antithetical to reproduction.

Weather station data shows that from Oct 5 to Oct 12, average humidity was 61.28% in College Station, Texas whereas from February 11 to 17 it was 69.62%, meaning these months were not drier. Instead, this period in February was on average 17.5 C colder than the trial in October, meaning to maintain temperatures of the growth chamber and breeding environment – especially during periodic freezes – HVAC systems were running more often, and thus lowered the *relative* humidity by expanding the volume and temperature of air.

| Table 6. Date ranges for experimental procedures, used to look up external weather conditions that may have affected ambient conditions of the rearing environment. | | |
| --- | --- | --- |
| Event | Trial 1 | Trial 2 |
| First Feeding | 2023-Sep-01 | 2023-Jan-12 |
| Second Feeding | 2023-Sep-08 | 2023-Jan-19 |
| Sift Prepupae | 2023-Sep-19 | 2023-Jan-26 |
| First Emergence | 2023-Oct-01 | 2023-Feb-05 |
| Sex and Sort into BugDorms | 2023-Oct-04 | 2023-Feb-09 |
| Load Cages (“Day 0”) | 2023-Oct-05 | 2023-Feb-11 |
| Start Observations (“Day 1”) | 2023-Oct-06 | 2023-Feb-12 |
| Box/Trap Added 1 Day After Peak Mating | 2023-Oct-08 | 2023-Feb-14* |
| First Eggs Harvested | 2023-Oct-08 | 2023-Feb-14 |
| End Observations | 2023-Oct-12 | 2023-Feb-15* |
| Final Eggs Harvested | 2023-Oct-12 | 2023-Feb-16* |
| Final Hatch Recording | 2023-Oct-18 | 2023-Feb-22 |

* Note: In Trial 1, the post-mating interval began on “Day 3”, but in trial 2 it began on “Day 2” or earlier for most flies; and so, attractant boxes/traps were added one day earlier for the latter. In Trial 2, Only eggs were harvested from Day 3.
